# Normalized level set model for segmentation of low-contrast objects in 2- and 3- dimensional images

**DOI:** 10.1101/2024.01.12.574651

**Authors:** Mirza M. Junaid Baig, Yao L. Wang, Samuel H. Chung, Armen Stepanyants

## Abstract

Analyses of biomedical images often rely on accurate segmentation of structures of interest. Traditional segmentation methods based on thresholding, watershed, fast marching, and level set perform well in high-contrast images containing structures of similar intensities. However, such methods can under-segment or miss entirely low-intensity objects on noisy backgrounds. Machine learning segmentation methods promise superior performance but require large training datasets of labeled images which are difficult to create, particularly in 3D. Here, we propose an algorithm based on the Local Binary Fitting (LBF) level set method, specifically designed to improve the segmentation of low-contrast structures.

## METHODS

### Normalized LBF model

The goal of image segmentation is to partition an image *I* : Ω → ℜ^*d*^ defined on a domain Ω ⊂ ℜ^*n*^ into foreground and background regions, making it possible to automatically detect and analyze the imaged objects. In the following, we will only consider grayscale images (*d* = 1) that are *n* = 2- and 3-dimensional, but the described method can be generalized on vector-valued images (e.g., color and hyperspectral) and images of higher dimensions (e.g., 3D plus time). The level set method [1] is ubiquitously used to solve segmentation problems. In a level set formulation [2-4], the segmenting contour *C* ⊂ Ω is defined as a zero-level of function *ϕ*: Ω → ℜ that defines the foreground as {*ϕ*(*x*) > 0| *x* ∈ Ω} and the background otherwise.

The level set function *ϕ* is determined in an optimization framework which usually combines image intensity-based costs and regularizing constraints on *ϕ*. For example, the Local Binary Field (LBF) level set model [2], minimizes a functional *E*_*LBF*_ (*ϕ, f*_1_, *f*_2_) which depends on the level set function *ϕ* and local average foreground and background intensity functions, *f*_1_ and *f*_2_.

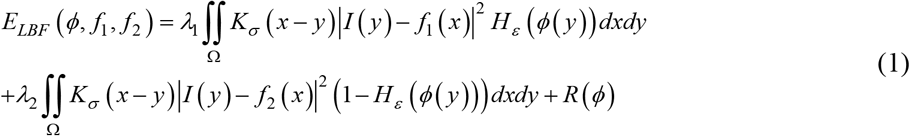

Here, the Gaussian kernel *K*_*σ*_ and the continuous approximation of the Heaviside step function *H*_*ε*_ are defined as follows:

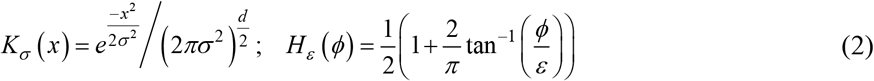

The first two terms in Eq. (1) represent the average squared deviations of image intensity from *f*_1_ and *f*_2_ in the foreground and background regions respectively, and the functional *R*(*ϕ*) can include any of the regularizing terms traditionally used with the level set method (see Eq. (4) for examples).

LBF segmentation works well in high-contrast images; however, when applied to images with stark intensity differences or high levels of noise, the method can miss or under-segment low-intensity structures. In an attempt to correct such problems, we introduced local normalizations in the foreground and background terms of the traditional LBF functional,

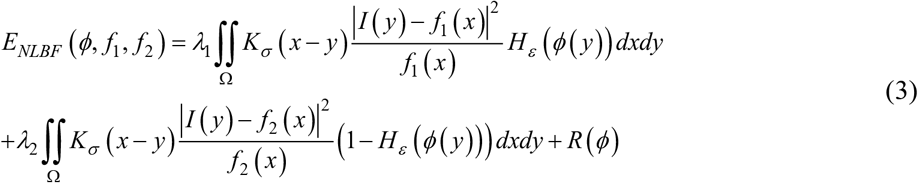

Direct comparisons of level set models published in the literature are hindered by the fact that they contain various regularizing terms, each included in the cost functional with a tunable parameter *µ*_*i*_ ≥ 0. To enable an unbiased comparison of the traditional and normalized LBF models, we considered 5 popular regularizers and systematically examined the corresponding parameter space focusing on sparse solutions in which 3 or more *µ*_*i*_ are zero.

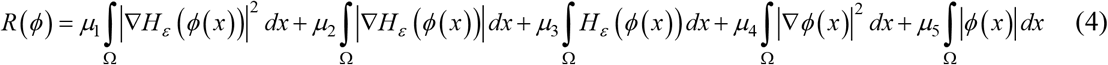

The second term in this expression corresponds to the length (or surface area) of the segmented contour *C* in *d* = 2 (or 3) dimensions respectively, while the third term represents the surface area (or volume) of the segmented objects. The other terms have no simple interpretation, but all prevent the level set function from diverging while permitting analytically tractable variational optimization of the cost functional,

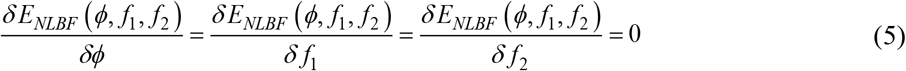

In these expressions, *δ* denotes variational derivative. Solution of Eqs. (5) leads to the following optimization algorithm based on gradient descent,

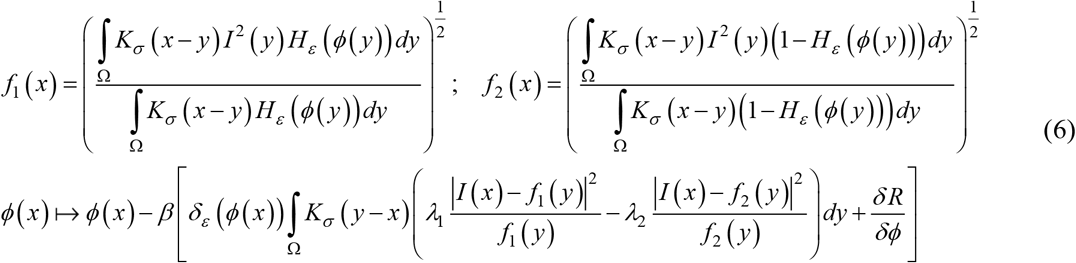

### Dataset and computational methods

The dataset used in this study (Figure 1) includes synthetic 2D and 3D images containing objects of distinct intensities and shapes with varying levels of added haze. These images were created in MATLAB and used to assess model performance, as their ground truth segmentations were known. In addition, the normalized LBF model was tested on 12 z-stacks of images of *C. elegans* neurons acquired with a Digital Micromirror Device (DMD) enhanced widefield microscope [5].

**Figure 1:**
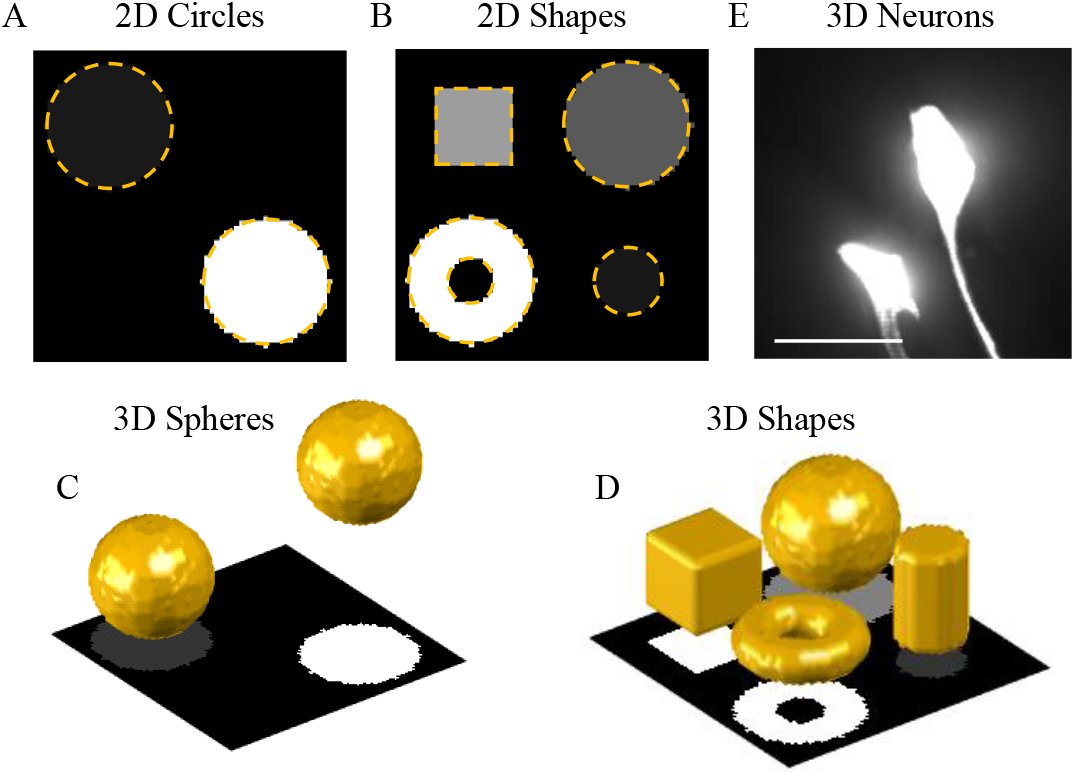
Dataset used for the analysis. **A**. Two circles of 1:10 intensity ratio with no added haze. Dashed lines outline the circles. 5 levels of haze were added to this image to create the 2D Circles images. **B**. Same for the 2D Shapes images. **C, D**. 3D images of different intensity structures. The 3D view (top) is shown together with the maximum intensity projection of the image (bottom). **E**. Maximum intensity projection of *C. elegans* neurons imaged in 3D with a DMD enhanced widefield microscope. Scale bar is 10 µm.

The synthetic 3D images were generated from two base images (Figure 1C, D). The first includes two equal size spheres of 1.0 and 0.1 intensities positioned at opposite corners of a 64×64×64 voxels image stack. The second image, 64×64×32 voxels in size, contains a torus, cube, cylinder, and sphere, of 1.0, 0.5, 0.25, and 0.10 intensities respectively. Haze was added to each base image at 5 levels of strength. Haze intensity was determined by blurring the base image with a Gaussian filter (isotropic standard deviation of 10 voxel sizes), rescaling the resulting image intensity to a maximum of 1, and, for each voxel, multiplying the result by a number chosen independently and uniformly at random from [‒*a, a*] interval. Five haze amplitudes *a* = 0.2, 0.3, 0.4, 0.5, and 0.6 were used, leading to 12 3D images in total. The 2D dataset was created by taking the maximum intensity projections of the two base 3D images (Figure 1A, B) and applying 5 levels of haze as described above, resulting in a total of 12 2D images. The ground truth segmentations for the 2D and 3D images contained zeros in the background and ones in the foreground.

The intensity- and continuity-related model parameters were fixed in all simulations with *λ*_1_ = *λ*_2_ =1, *σ* = 10, and *ε* = 1. The optimal values of parameters *µ*_1-5_ were determined for each model and image type independently with a grid search algorithm. Each model was run for 250 iteration steps or until the relative change in the cost functional dropped below 10^-6^, whichever came first. Intersection Over Union (IOU) of model results and the corresponding ground truths were used to access the performance.

## RESULTS

Figure 2 illustrates the process of segmentation for the traditional and normalized versions of the LBF algorithms. Normalized LBF (Figure 2C) could segment the low-intensity cylinder with greater accuracy, producing a smoother surface that resembles the ground truth more than that generated by traditional LBF (Figure 2B).

**Figure 2:**
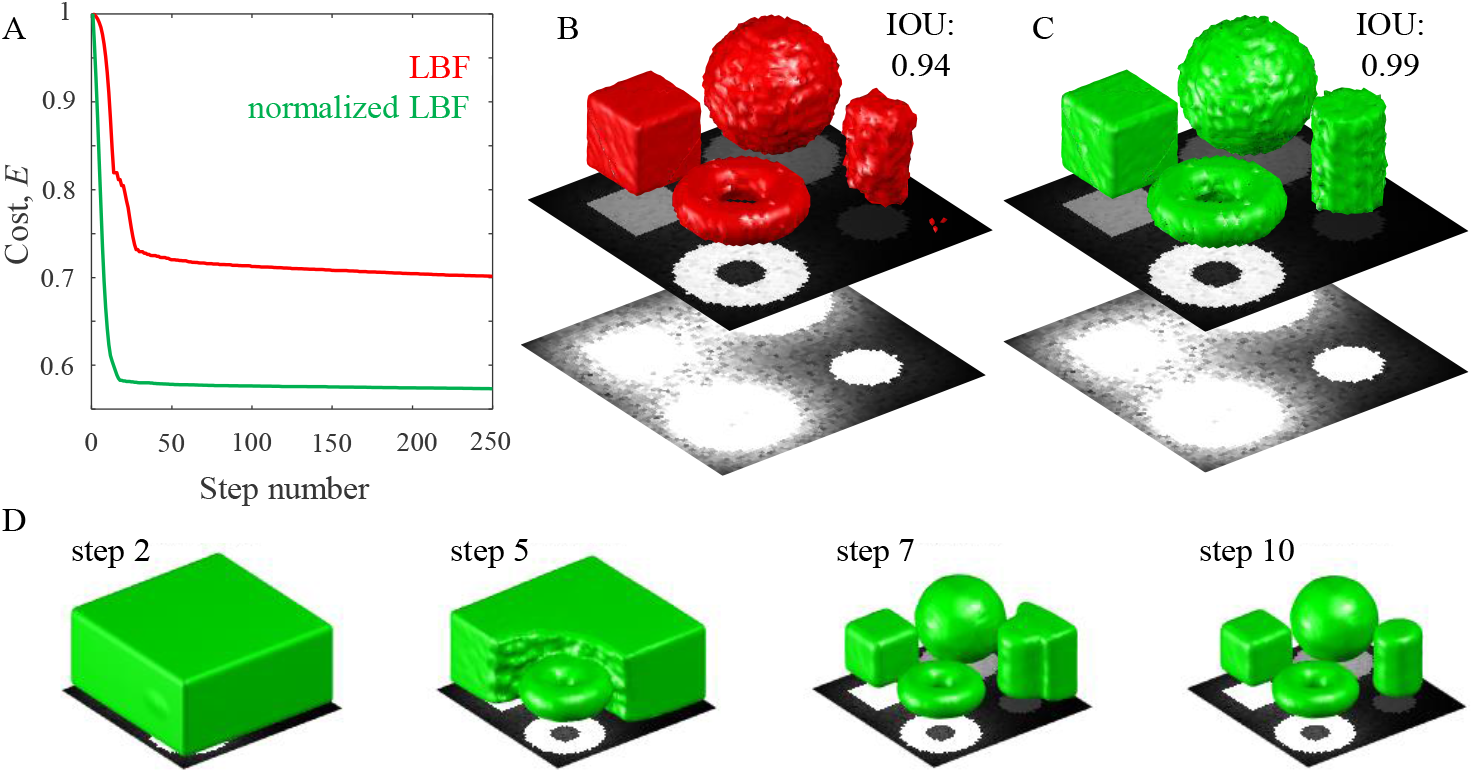
Performance of the traditional (red) and normalized LBF (green) models. **A**. Normalized cost functionals vs. number of iteration steps. **B**. Segmentation results of the LBF model overlayed on the maximum intensity projection (top plane) of the 3D image. Same image at 10× saturation (bottom plane) shows hard to see shapes and haze (*a* = 0.2). **C**. Same for the normalized LBF model. **D**. The normalized LBF model typically converges to an adequate segmentation within ∼10 iteration steps.

To substantiate this result, both models were tested on all the synthetic 2D and 3D images. To enable an unbiased comparison, a single set of best parameters was determined for each model (see Methods) in the 2D and 3D cases. The normalized LBF model yields significantly higher IOU scores than traditional LBF (Table 1).

**Table 1:** IOU (mean ± SE) for 2D and 3D segmentation results of the traditional and normalized LBF models (***p<0.001, unequal variance *t*-test).

|  | LBF | Normalized LBF |
| --- | --- | --- |
| 2D images ($n = 12$) | $0.85 \pm 0.01$ | $0.94 \pm 0.02^{***}$ |
| 3D images ($n = 12$) | $0.92 \pm 0.02$ | $0.99 \pm 0.01^{***}$ |

The normalized LBF method was also tested on the segmentation of *C. elegans* neurons that were fluorescently labeled and imaged in 3D with a widefield microscope. Figure 3 shows examples of results for the ASJ and AIY neuron types. Visual inspection of these results demonstrates the capability of the method to accurately delineate neuron structures in complex biological images.

**Figure 3:**
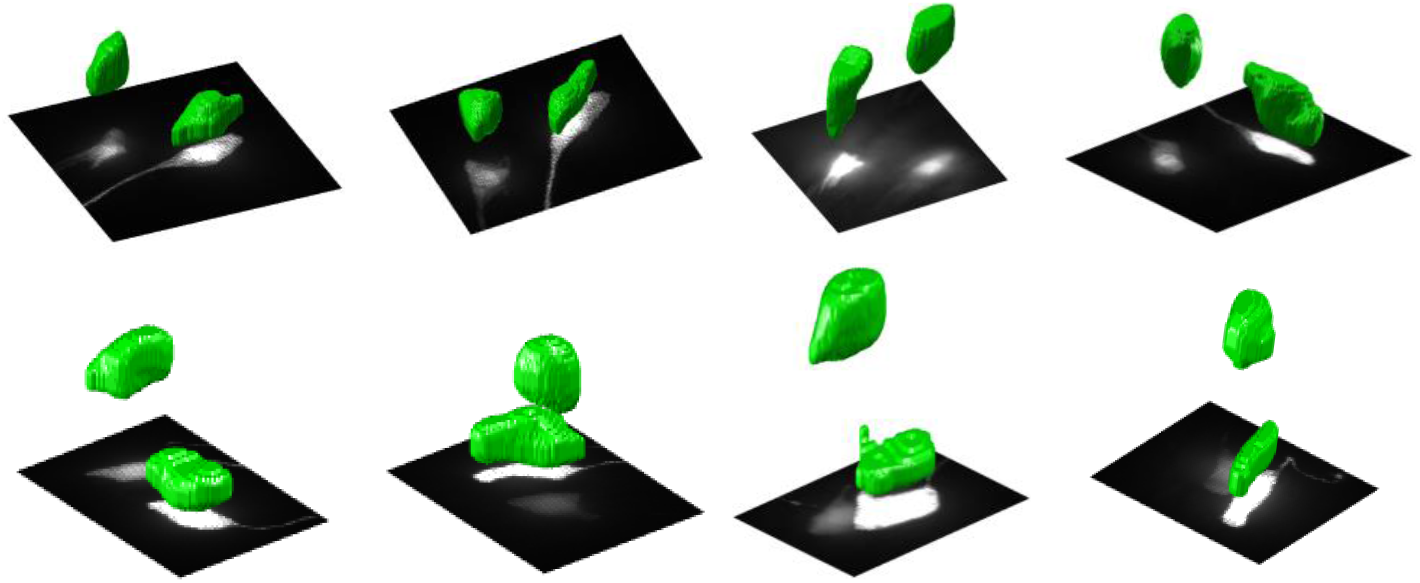
Normalized LBF model can be used to segment cell bodies of *C. elegans* neurons despite significant differences in their contrast and the surrounding haze. **Top row**. 3D segmentation of AIY neuron cell bodies (green) shown over maximum intensity projections of image stacks. **Bottom row**. Same for the ASJ neurons.

## CONCLUSIONS

We propose a new segmentation method based on the LBF level set, designed to improve the segmentation of biomedical images containing structures of varying intensities and contrasts. The method was tested on a synthetic dataset of 2D and 3D images with known ground truths, showing that the proposed normalization in the traditional LBF model improves its performance without jeopardizing computational tractability. The method was also used to segment the cell bodies of *C. elegans* neurons in 3D stacks of fluorescent images, where all neurons could be accurately segmented despite their large intensity differences and the surrounding haze.

This work was supported by the NIH grant R56 NS128413. The code and the dataset of images used in this study are available at https://github.com/neurogeometry/Normalized_Level_Set.

